# Mitochondrial LonP1 protease is implicated in the degradation of unstable Parkinson disease-associated DJ-1/PARK 7 missense mutants

**DOI:** 10.1101/2020.09.29.318683

**Authors:** Raúl Sánchez-Lanzas, José G. Castaño

**Affiliations:** Departamento de Bioquímica, Instituto de Investigaciones Biomédicas “Alberto Sols”, UAM-CSIC. Facultad de Medicina UAM. 28029 Madrid. Spain

**Keywords:** DJ-1/PARK7, Parkinson, proteolysis, mitochondria, LonP1, proteasome, autophagy, lysosome

## Abstract

*DJ-1/PARK7* mutations are linked with familial forms of early onset Parkinson disease (PD). We have studied the degradation of untagged DJ-1 WT and missense mutants in mouse embryomic fibroblasts obtained from DJ-1 null mice, an approach closer to the situation in patients carrying homozygous mutations. The results showed that the mutants: L10P, M26I, A107P, P158DEL, L166P, E163K and L172Q are unstable proteins, while A39S, E64D, R98Q, A104T, D149A, A171S, K175E and A179T are as stable as the DJ-1 WT. Inhibition of proteasomal and autophagic-lysosomal pathways had little effect on their degradation. Immunofluorescence and biochemical fractionation studies indicated that M26I, A107P, P158DEL, L166P, E163K and L172Q mutants associate with mitochondria. Silencing of mitochondrial matrix protease LonP1 produced a strong reduction of the degradation of those mitochondrialy associated DJ-1 mutants, but not of mutant L10P. These results demonstrated a mitochondrial pathway of degradation of those DJ-1 missense mutants implicated in PD pathogenesis.

## Introduction

The majority of Parkinson’s disease (PD) patients are sporadic cases; rare familiar forms of PD can account for 5-10% of all PD cases. Mutations in several genes are well established as a genetic cause in familial PD (Hernandez et al., 2016). PARK7 /DJ-1 is one of those PD-linked genes whose pathogenic mutations show autosomal recessive inheritance and early-onset of the PD phenotype (Bonifati et al., 2003). Those genetic mutations include CNVs (exonic deletions and truncations), splice-site mutations, homozygous (L10P, M26I, E64D, P158DEL, E163K, L166P and L172Q) and heterozygous (A39S, A104T and D149A) missense mutations, and rare polymorphisms (R98Q, A171S) in healthy individuals that are not associated with PD (Nuytemans et al., 2010) (Hernandez et al., 2016).

Native DJ-1 protein is a dimer with a flavodoxin-like helix-strand-helix structure (Tao and Tong, 2003; Honbou et al., 2003; Wilson et al., 2003) (Huai et al., 2003; Lee et al., 2003) (Malgieri and Eliezer, 2008). DJ-1, initially identified as a potential oncogene cooperating with Ha-Ras in cell transformation (Nagakubo et al., 1997), is implicated in several pathways, such as: transcriptional regulation (Takahashi et al., 2001) (Taira et al., 2004a) (Xu et al., 2005) (Shinbo et al., 2005) (Zhong et al., 2006), RNA binding (Hod et al., 1999) (van der Brug et al., 2008), regulation of sumoylation (Shinbo et al., 2006), protein folding as a chaperon (Shendelman et al., 2004) (Zhou et al., 2006) (Deeg et al., 2010) or co-chaperon (Lee et al., 2018) and cell death (Junn et al., 2005) (Tai-Nagara et al., 2014). DJ-1 protein is cytoprotective being a sensor of oxidative stress and acting as antioxidant preventing apoptosis (Meulener et al., 2006) (Canet-Aviles et al., 2004) (Taira et al., 2004b) (Zhou and Freed, 2005) (Clements et al., 2006) (Aleyasin et al., 2007) (Liu et al., 2008; Blackinton et al., 2009) (Aleyasin et al., 2010) (Billia et al., 2013) (Choi et al., 2014). Taking into account that some *PARK7/DJ-1* gene mutations result in a loss of function of the gene (no protein produced), it is reasonable to hypothesize that also some of the missense point mutants may produce a loss of function of the protein. In agreement with this hypothesis, accelerated protein degradation (increased protein instability in the cell) of the DJ-1 point mutants would produce a decrease in the steady-state levels of the protein mimicking, in part, the phenotype of loss of gene function (no protein produced). In fact, DJ-1 L166P mutant has reduced stability in the cell (Bonifati et al., 2003) (Macedo et al., 2003) (Miller et al., 2003) (Moore et al., 2003) (Olzmann et al., 2004) (Takahashi-Niki et al., 2004) (Blackinton et al., 2005) (Gorner et al., 2007) (Alvarez-Castelao et al., 2012a). DJ-1 M26I mutant have increased degradation in the cell according to some reports (Takahashi-Niki et al., 2004; Blackinton et al., 2005), while other groups found no effect of the M26I mutation on protein stability (Moore et al., 2003) (Baulac et al., 2004) (Xu et al., 2005) (Alvarez-Castelao et al., 2012a). The mutants A104T and D149A have also increased rates of degradation (Blackinton et al., 2005), but found stable by other groups (Moore et al., 2003) (Baulac et al., 2004) (Zhang et al., 2005; Xu et al., 2005) (Alvarez-Castelao et al., 2012a). More recently described missense mutants of DJ-1 L10P, P158DEL and L172Q are also unstable proteins (Ramsey and Giasson, 2010) (Rannikko et al., 2012) (Repici et al., 2013) (Taipa et al., 2016). Finally, the point mutant E163K retains similar properties to DJ-1 wild type (WT) protein respect to stability in cells (Ramsey and Giasson, 2008), but it reduces the thermal stability of DJ-1 in solution disrupting a salt bridge of E163 with R145 (Lakshminarasimhan et al., 2008).

Several studies of the degradation of DJ-1 missense mutants use tagged versions of the mutants and transfection into recipient cells expressing their endogenous wild-type DJ-1., From our point of view the use of tagged constructs is inadequate to study protein degradation. When a protein is tagged either in its N-terminus or its C-terminus it is assumed that the behaviour of this protein in the cell will be equivalent to that of the untagged endogenous protein, but this assumption is not necessarily true. As a proof of the previous statement, N-terminal tagged DJ-1 L166P has a reduced degradation rate compared to untagged DJ-1 L166P, while tagging L166P at the C-terminus has less effect (Alvarez-Castelao et al., 2012b). Another caveat to correctly interpreted the results reported is that the DJ-1 missense mutants variably heterodimerize with the endogenous DJ-1 WT of the recipient cells, and those interactions may also, in principle, modify the stability of the mutant protein. This situation will not occur in patient cells where only the missense mutant DJ-1 protein is expressed (homozygous mutation). A way to circumvent those caveats is the study of the stability of the untagged DJ-1 missense mutants in null DJ-1 cells. The use of null DJ-1 mouse embryonic fibroblasts (MEFs) to study DJ-1 missense mutants stability have been used in some reports (Rannikko et al., 2012) (Repici et al., 2013). Here we have systematically investigated the degradation of untagged human DJ-1 wild type and its missense point variants: L10P (Guo et al., 2008), M26I (Abou-Sleiman et al., 2003), A39S (Tang et al., 2006), E64D (Hering et al., 2004), R98Q (Abou-Sleiman et al., 2003), A104T (Clark et al., 2004), A107P (Macedo et al., 2009), D149A (Abou-Sleiman et al., 2003), P158DEL (Macedo et al., 2009), E163K (Annesi et al., 2005), A171S (Clark et al., 2004), L172Q (Taipa et al., 2016), K175E (Nuytemans et al., 2009) and A179T (Nuytemans et al., 2009; Macedo et al., 2009) in MEFs from null DJ-1 mice. The results have shown that DJ-1 pathogenic point mutants; L10P, M26I, A107P, P158DEL, E163K, L166P and L172Q showed a significant increase of the degradation rate respect to DJ-1 WT in MEFs from null DJ-1 mice, and no significant difference respect to WT was found for A39S, E64D, A104T, D149A, K175E, A179T missense mutants or the rare R98Q, A171S polymorphic variants.

## Results

### Degradation of human DJ-1 wild type and missense mutants

As previously stated in the introduction, degradation of ectopically expressed DJ-1 missense mutants could be affected by interactions with the cell endogenous DJ-1 and/or the use of tagged constructs Accordingly, untagged DJ-1 wild type (WT) and the missense mutants were transfected into null DJ-1 mouse embryonic fibroblasts (MEF). The mutants: L10P, M26I, A107P, P158DEL, E163K, L166P and L178Q were unstable (Fig. 1 A and B) after treatment of cells with cycloheximide (CHX), and showed different degradation rates. In contrast, DJ-1 WT and the missense mutants A39S, E64D, R98Q, A104T, D149A, A171S, K175E and A179T were not significantly degraded after treatment of the cells for 24h with CHX (Fig. 2). The apparent degradation rate of DJ-1 missense mutants, evaluated by the corresponding half-life, and ordered from smallest to longest half-life, was: L10P, P158DEL and L166P < A107P < L172Q < E163K < M26I (Supplementary Table I)). We have previously shown that transfected human DJ-1 WT and point mutants M26I, R98Q, A104T, D149A are stable proteins in N2a mouse cells (expressing the endogenous mouse DJ-1), and L166P is unstable (Alvarez-Castelao et al., 2012a). We extended those previous studies with similar assays to other missense DJ-1 mutants in the same cell line. In those experiments, the protein levels of the missense mutants: A39S, A171S, K175E and A179T did not significantly change after treatment of transfected cells with CHX for 24h (Supplementary Fig. 1A). In contrast, the point mutants L10P, A107P, P158DEL, E163K, L166P (shown again for comparison) and L172Q were degraded at different rates (Supplementary Fig. 1B).

**Legend to Fig. 1.**
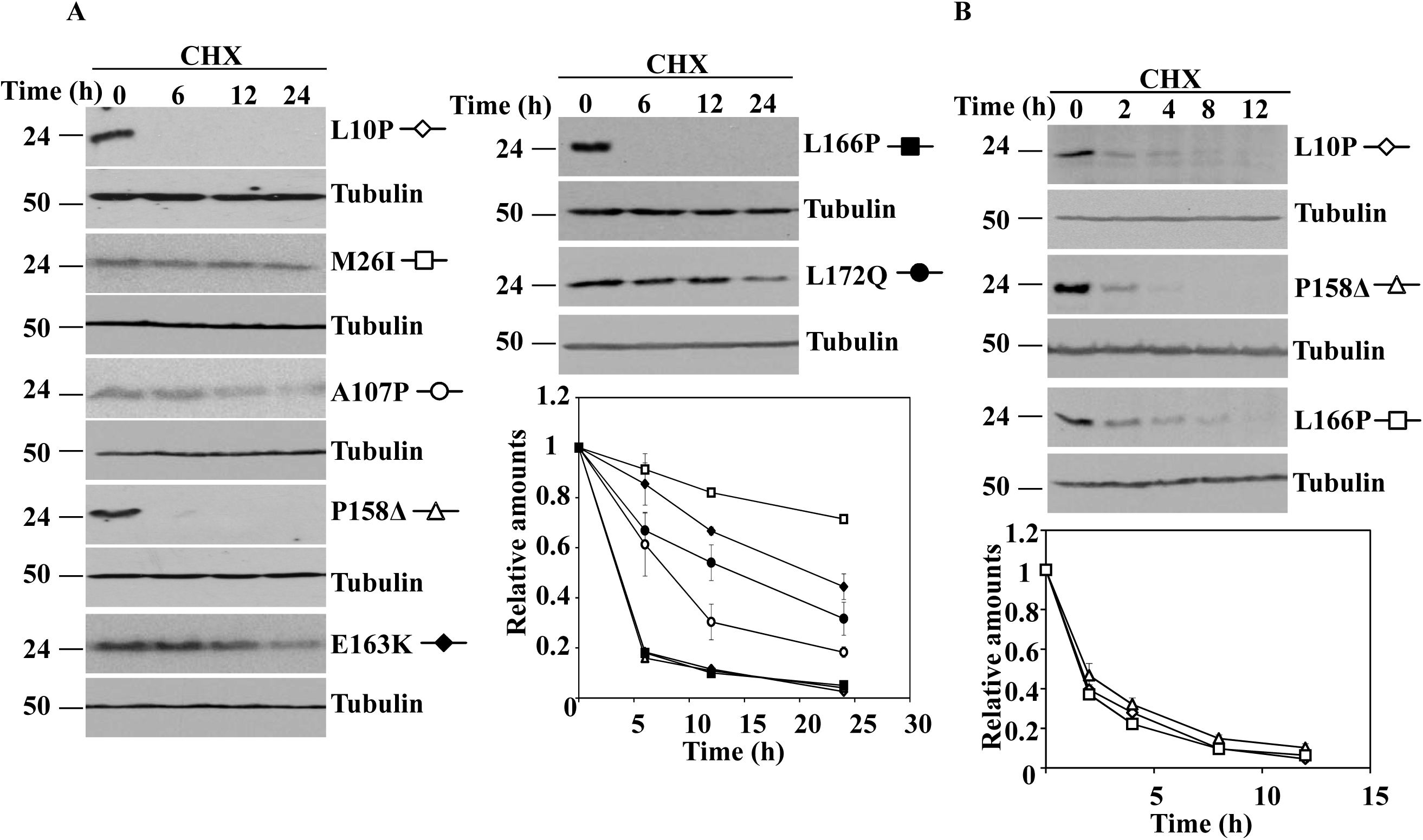
Time-course of the degradation of human DJ-1 unstable missense mutants transfected in DJ-1 null MEFs. DJ-1 null MEFs were transiently transfected with the indicated untagged human DJ-1 (hDJ-1) constructs and 48 hours after transfection were treated with 25µg/mL cycloheximide (CHX) for the times indicated. A, panels show representative immunoblots probed with anti-DJ-1 polyclonal antibodies of cells transfected with hDJ-1 missense mutants L10P, M26I, A107P, P158Δ, E163K, L166P and L172Q. B, panels show the results obtained with transfected DJ-1 missense mutants L10P, P158Δ and L166P in cells treated with CHX for shorter times, as indicated. Anti-tubulin antibodies were used as total protein loading control. Quantifications from three different experiments are shown in the graphs as mean ± s.e.m.

**Legend to Fig. 2.**
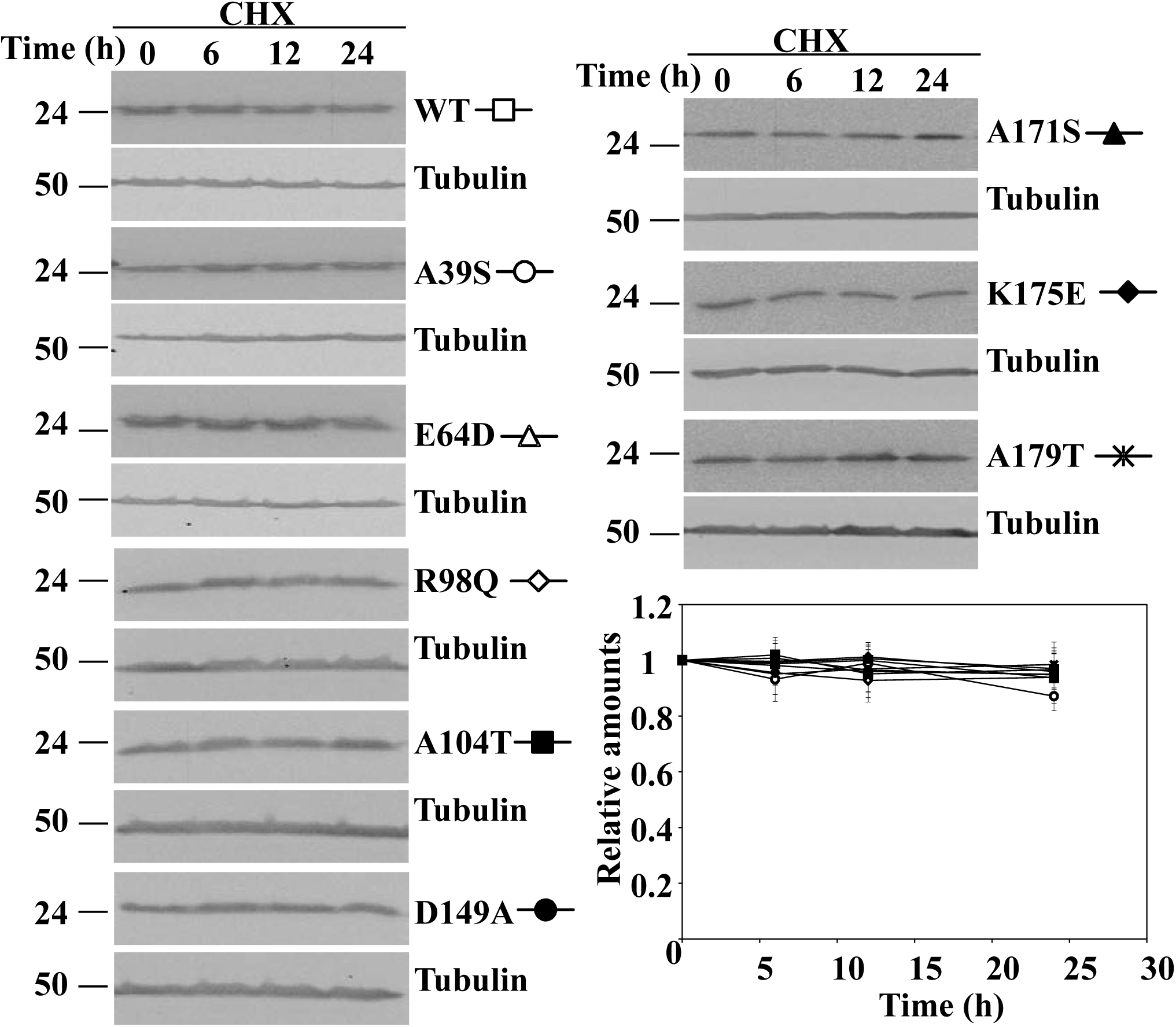
Degradation of human DJ-1 wild type and missense mutants in transfected DJ-1 null MEFs. DJ-1 null MEFs were transiently transfected with the indicated untagged human DJ-1 (hDJ-1) constructs and 48 hours after transfection were treated with 25µg/mL cycloheximide (CHX) for the times indicated. Panels show representative immunoblots developed with anti-DJ-1 polyclonal antibodies of hDJ-1 wild type (WT), A39S, E64D, R98Q, A104T, D149A, A171S, K175E and A179T. Anti-tubulin antibodies were used as total protein loading control. Below is shown the graph of quantification of the corresponding immunoblots. Data are mean ± s.e.m from three different experiments.

Although the proteasomal pathway is implicated in the degradation of DJ-L166P, its degradation is only partially prevented by proteasome inhibitors (Alvarez-Castelao et al., 2012a),. As a consequence, we decided to re-evaluate those results and to test if inhibition of other proteolytic pathways can be more effective. To that end, transfected cells were treated with CHX for 24h for those missense mutants that have shown longer half-lives M26I, A107P; E163K and L172Q and for 12h for those missense mutants with shorter half-lives L10P, P158DEL and L166P, in the absence or in the presence of several protease inhibitors. As shown in Fig. 3, addition of MG132 (proteasome inhibitor) or inhibitors of the autophagic-lyssosomal pathway (NH_4_Cl, NH_4_Cl in combination with leupeptin or 3-methyl adenine together with E64) had no significant effect on the degradation of the missense mutants (or DJ-1 WT). Those results suggested that neither the proteasomal nor the autophagic-lysosomal pathways of protein degradation played a major role in the degradation of the unstable DJ-1 missense mutants.

**Legend to Fig. 3.**
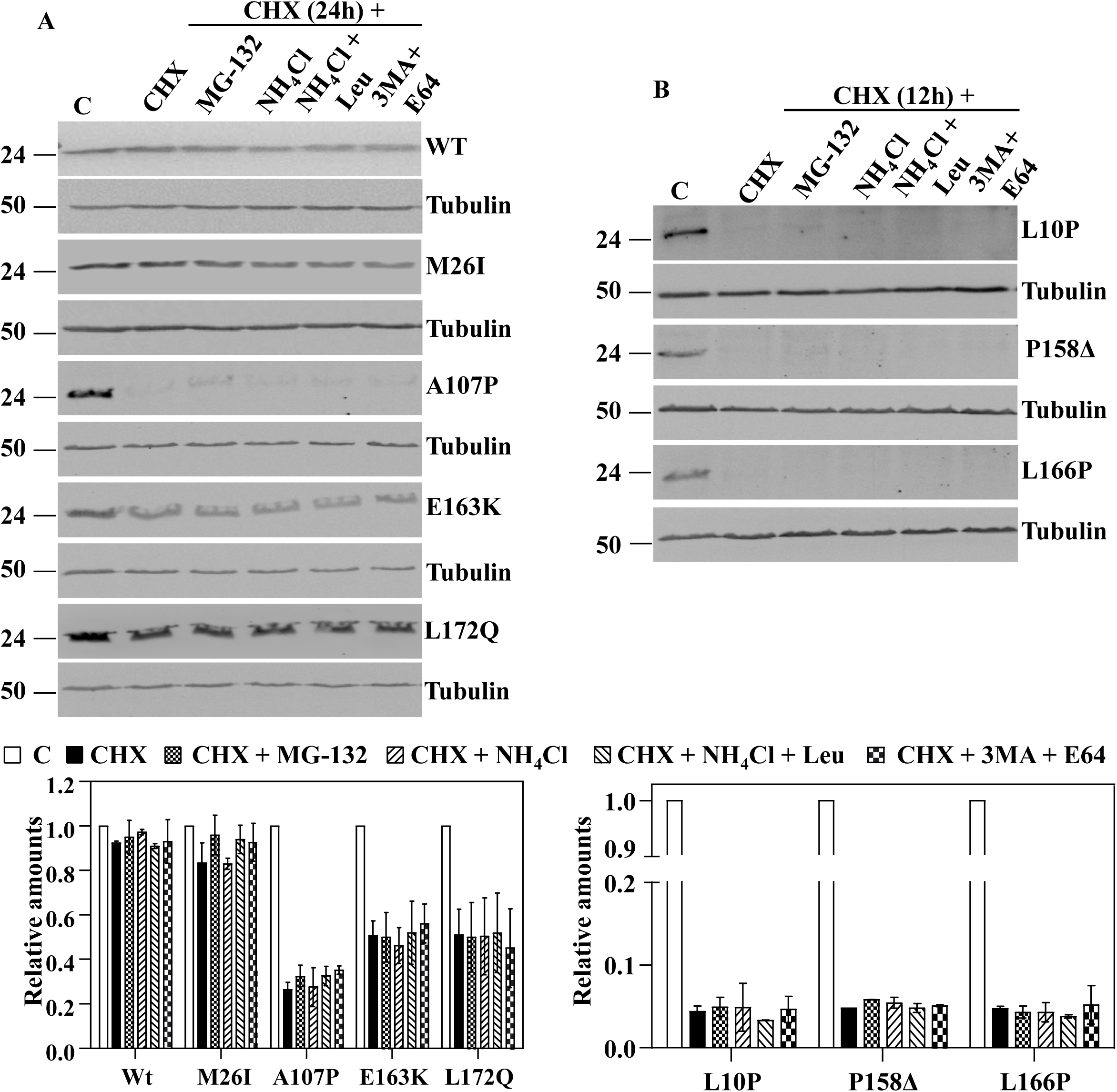
Effect of protease inhibitors on the degradation of human DJ-1 wild type and missense mutants transfected in DJ-1 null MEFs. DJ-1 null MEFs were transiently transfected with the indicated untagged human DJ-1 (hDJ-1) constructs and 48 hours after transfection were kept in complete medium (C) or treated with 25µg/mL cycloheximide (CHX) in the absence (DMSO) or the presence of 10µM MG-132, 20mM NH_4_Cl, 20mM NH_4_Cl plus 50µM leupeptin (Leu) or 10mM 3-methyl adenine (3MA) plus 5µM E64 for 12 or 24 hours, as indicated. Total cell lysates were analysed by Western and immunoblot with the corresponding specific antibodies. A, panels show representative immunoblots with anti-DJ-1 polyclonal antibody of hDJ-1 wild type (WT), M26I, A107P, E163K and L172Q. B, panels show the results obtained with transfected hDJ-1 L10P, P158Δ and L166P developed with anti-DJ-1 polyclonal antibody. Anti-tubulin antibodies were used as total protein loading control. Graphs below each panel show the quantifications of the levels of DJ-1 protein respect to untreated cells as control. Values are expressed as mean ± s.e.m from three different experiments.

### Subcellular localization of human DJ-1 wild type and missense mutants

A possible clue to understand the pathway of degradation of DJ-1 missense mutants could be their subcellular localization. DJ-1protein is described to be localized both in the cytoplasm and in the nucleus (Nagakubo et al., 1997) (Bonifati et al., 2003) (Bandyopadhyay and Cookson, 2004) and to translocate to the mitochondria upon cell oxidative stress (Zhang et al., 2005) (Li et al., 2005). L166P missense mutant is reported to be localized in the cytoplasm and mitochondria (Bonifati et al., 2003) (Kojima et al., 2016), E163K mutant was shown to be localized in the mitochondria (Bjorkblom et al., 2014a) (Kojima et al., 2016), as well as M26I (Kojima et al., 2016) and L172Q (Taipa et al., 2016) missense mutants. Therefore, the subcellular localization of the human DJ-1 WT and missense mutants was determined after transfection of DJ-1 null MEF by indirect immunofluorescence. Fig. 4 shows the results obtained and Supplementary Fig.2 shows the analysis of the co-localization of DJ-1 and mitotracker fluorescences. Without doubt, it can be concluded that the immunofluorescence signal of M26I, A107P, P158DEL, E163K, L166P and L172Q co-localize with mitotracker fluorescence (Pearson coefficient ≥. 0.6).

**Legend to Fig. 4.**
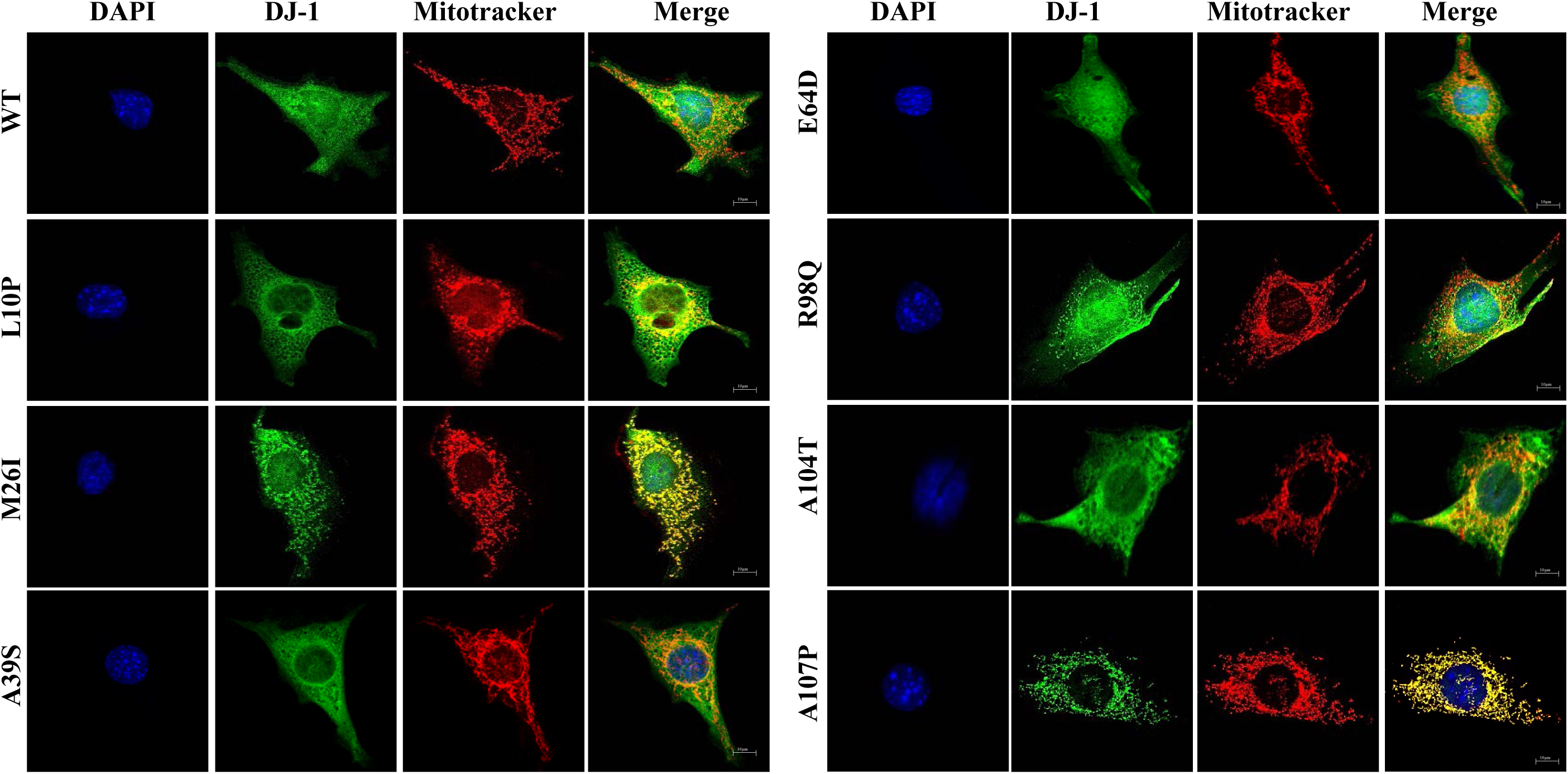
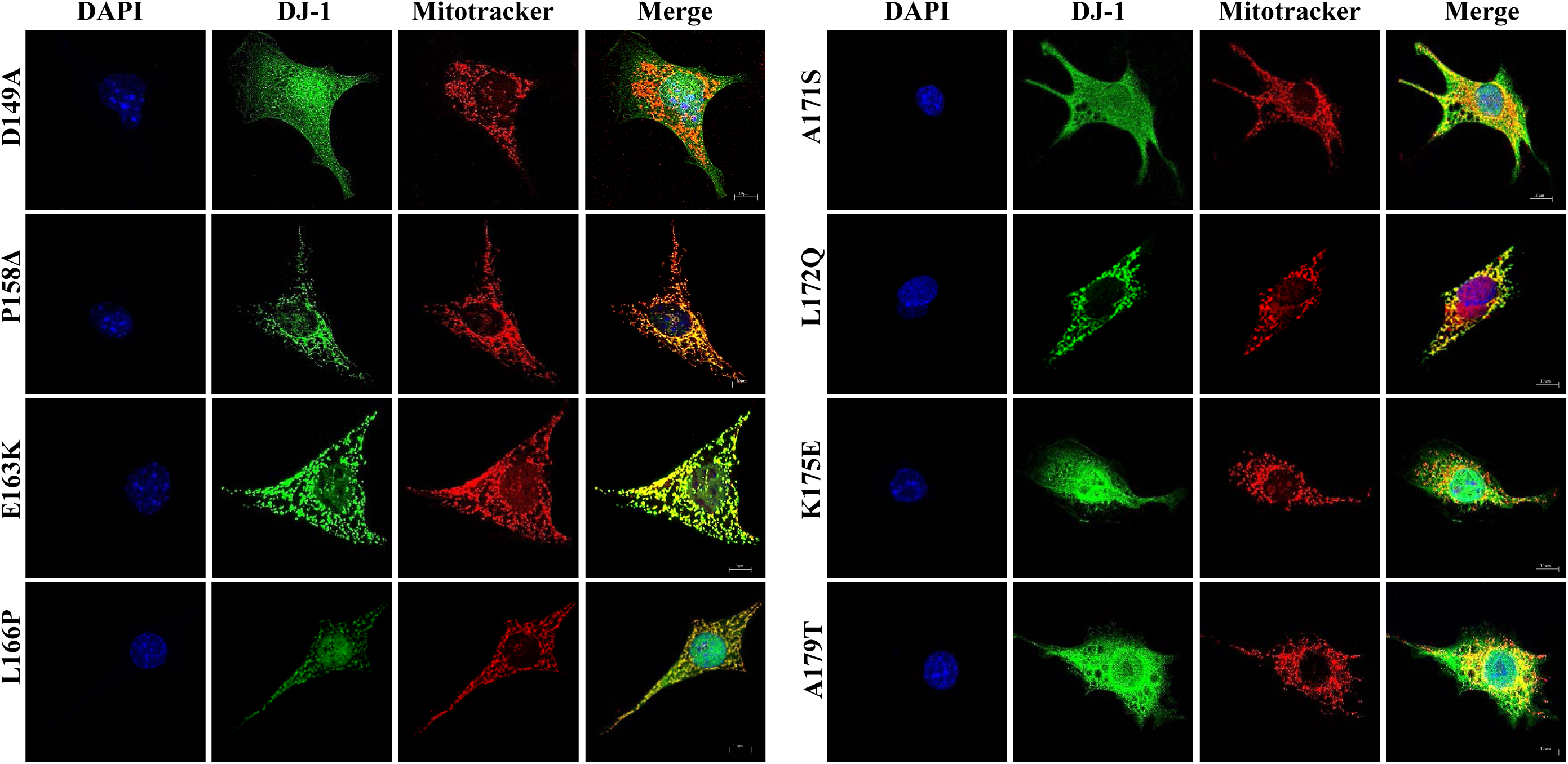
Subcellular localization by indirect immunofluorescence of human DJ-1 wild type and missense mutants transfected in DJ-1 null MEFs. Confocal fluorescence images of the indicated untagged human DJ-1 constructs in transfected DJ-1 null MEFs grown under basal conditions (complete medium), stained with Mitotracker (red), processed for immunofluorescence with anti-DJ-1 polyclonal specific antibodies (green) and counterstained with DAPI for nuclei visualization (blue). Quantifications of the co-localization of red (mitochondria) and green (DJ-1 immunofluorescence) pixels are presented in Supplementary Figure 2.

In contrast, a nuclear cytoplasmic fluorescence with some co-localization (naked-eye observation, Pearson coefficient >0.2 and <0.4) with mitotracker was observed for E64D, R98Q, A104T, D149A and A171S missense mutants. Finally, nuclear and cytoplasmic distribution of fluorescence without significant co-localization with mitotracker fluorescence (Pearson coefficient ≤0.2) was found for DJ-1 WT, L10P, A39S, A171S and K175E. These results suggested that a good approach to study both, protein degradation and subcellular localization, would be to obtain fusion constructs of DJ-1 mutants with fluorescent proteins. To that end, constructs of M26I and L166P fused to the N-terminus of EGFP were produced. The results obtained after transfection in MEFs from null DJ-1 mice (see Supplementary Fig. 3) clearly indicated that the C-terminal tagging of the unstable L166P missense mutant greatly slows down its degradation rate compared to the untagged version (see Fig. 1 for comparison), similarly M26I fusion construct was not significantly degraded after 24h of incubation in the presence of CHX. Furthermore, the fusion proteins did not reproduce the preferential mitochondrial localization observed by indirect immunofluorescence of the untagged M26I and L166P DJ-1 missense mutants (compare Supplementary Fig. 3 and Fig. 4). In conclusion, the approach with fluorescent fusion constructs to study protein degradation and subcellular localization was discarded. To get independent evidence of the subcellular localization, biochemical fractionations studies of DJ-1 mutants transfected in DJ-1 null MEFs were performed. Fig. 5 shows the results obtained. Cell fractionation studies showed that DJ-1 point mutants M26I, A107P; P158DEL, L166P, E163K and L172Q showed significant association with the mitochondrial fraction, while they were also clearly present in the cytoplasmic fraction. This interpretation is further strengthened by comparison to similar fractionation studies with transfected DJ-1 WT and missense point mutants L10P, A39S, E64D, R98Q, A104T, D149A, A171S, K175E and A179T that showed a predominant cytoplasmic distribution.

**Legend to Fig. 5.**
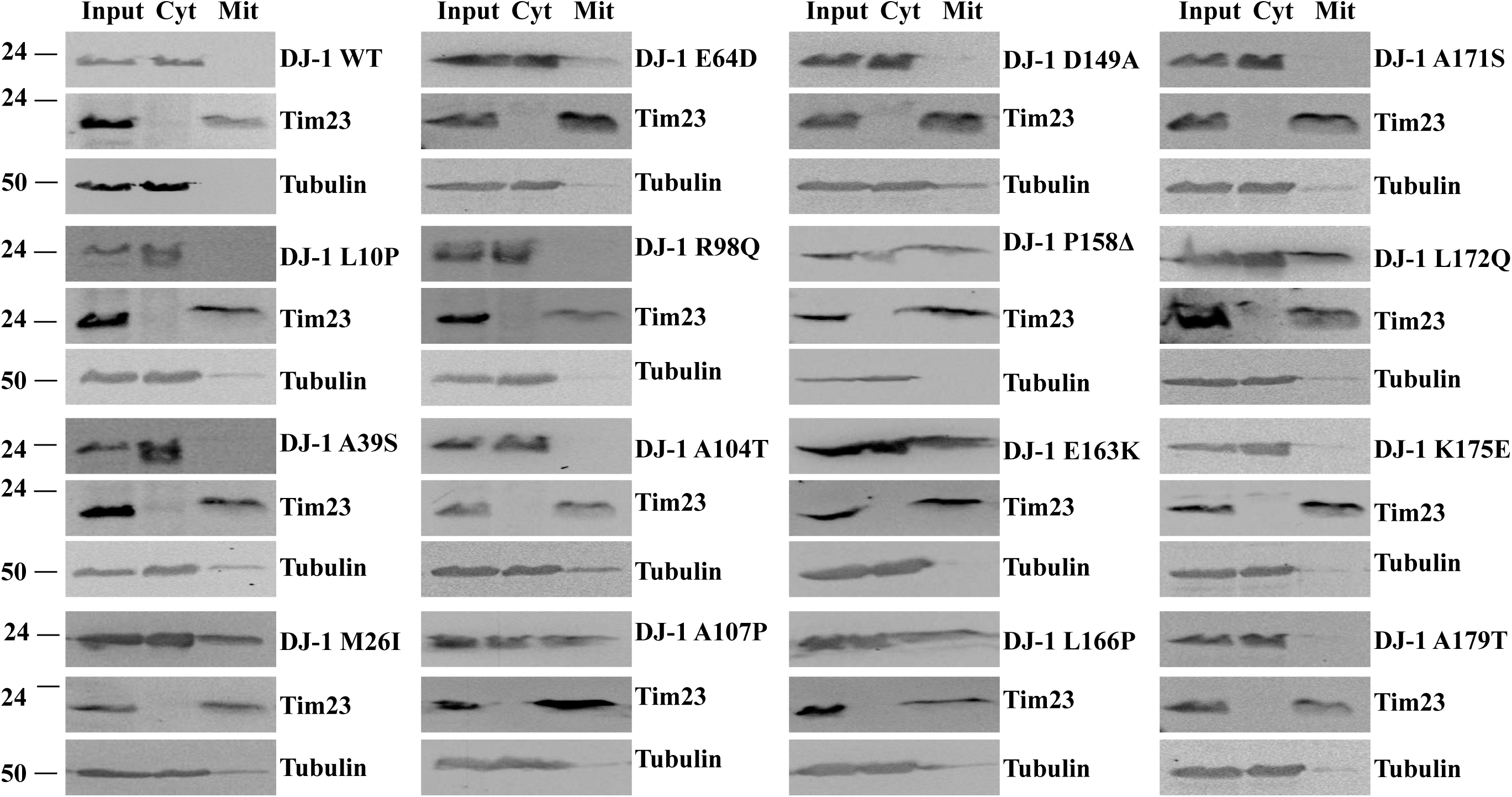
Biochemical cell fractionation studies of human DJ-1 wild type and missense mutants transfected in DJ-1 null MEFs. DJ-1 null MEFs were transiently transfected with the indicated untagged human DJ-1 constructs and 48h after transfection were processed for subcellular fractionation, as described under the Material and Methods section. Proteins from whole cell lysates (Input), cytoplasmic fraction (Cyt) and mitochondrial fraction (Mit) were analysed by Western and immunoblot with the indicated specific antibodies: anti-DJ-1 polyclonal antibody, anti-Tim23 as a mitochondrial marker and anti-tubulin as a cytoplasmic marker. Panels show representative immunoblots from two different experiments.

### Pathway of degradation of DJ-1 unstable missense mutants

The above results seemed to indicate that a mitochondrial pathway could be responsible of the degradation of the unstable DJ-1 mutants. The mitochondrial localization of some unstable DJ-1 mutants and some previous reports describing that DJ-1 may translocate to the mitochondria matrix (Bjorkblom et al., 2014b; Kojima et al., 2016) prompt us to study the possible involvement of mitochondrial matrix proteases in the degradation of DJ-1 missense mutants. The matrix mitochondrial LonP1 protease was examined as a possible candidate. LonP1 is an essential gene in mice (Quiros et al., 2014). as consequence we were unable to get cells from LonP1 null mice. Consequently, shRNA interference of mouse LonP1 (mLonP1) in null DJ-1 MEF or N2a cells were used to test this hypothesis. As shown in Fig. 6, the down regulation of LonP1 expression in null DJ-1 MEFs (Fig. 6A immunoblot and Fig.6 B immunofluorescence) resulted in a significant decrease in the degradation rate of unstable DJ-1 missense mutants A107P, P158DELTA, E163K, L166P and L172Q (Fig. 6 C and D), but not of DJ-1 L0P. Similar results were obtained with mLonP1 silencing in N2a cells expressing those missense DJ-1 mutants (Supplementary Fig. 4).

**Legend to Fig. 6.**
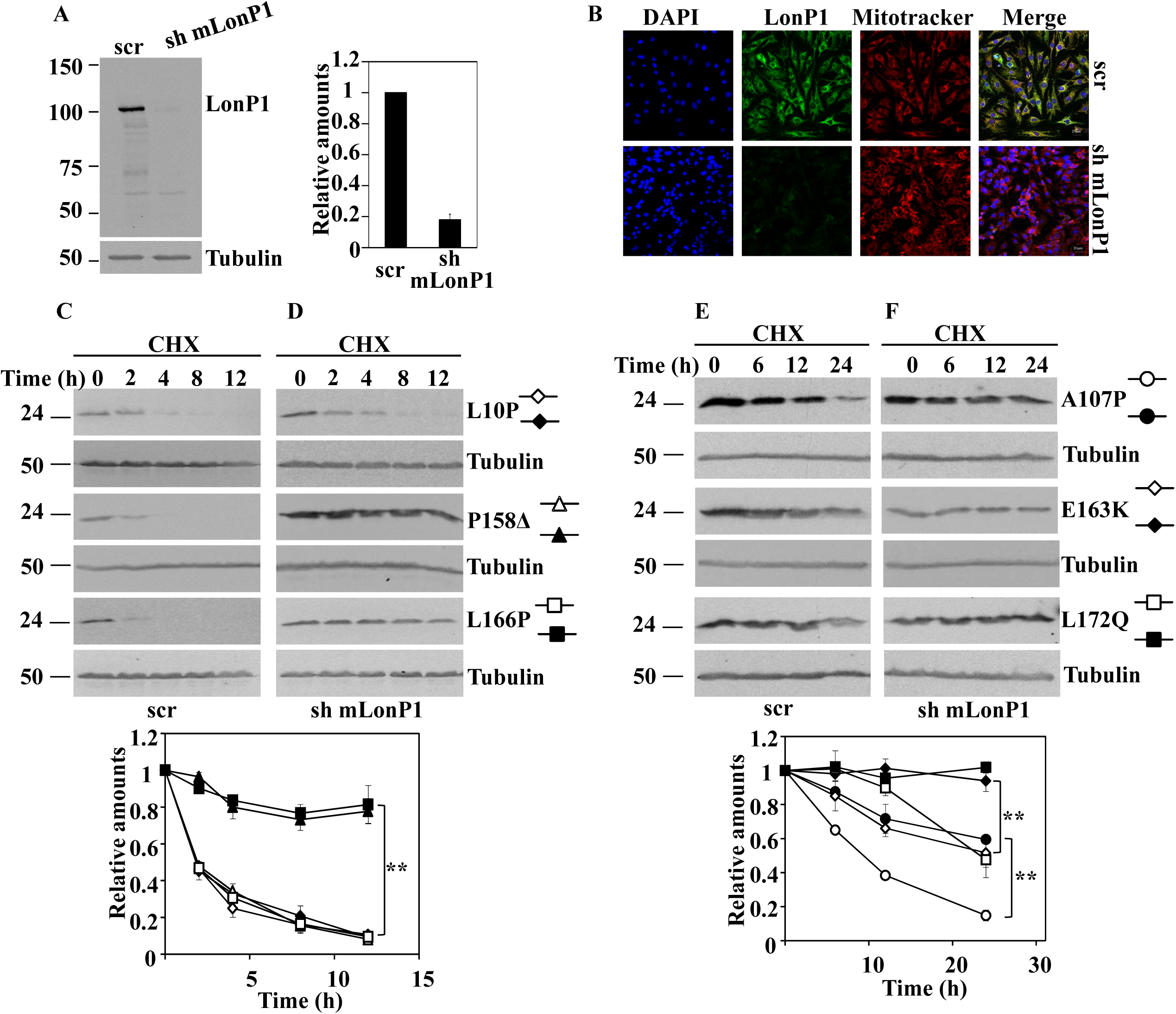
Effect of LonP1 silencing on the degradation of DJ-1 unstable mutants transfected in DJ-1 null MEFs. DJ-1 null MEFs were transduced with either scrambled shRNA (scr) or specific shRNA for mouse LonP1 (sh mLonP1), as described under the Material and Methods section. A, panel shows a representative immunoblot developed with specific anti-LonP1 antibodies of LonP1 shRNA-mediated knockdown in DJ-1 null MEFs. Quantification is shown in the right graph. B, confocal fluorescence images of non-target shRNA lentiviral transduced (scr) or LonP1 shRNA lentiviral transduced (sh mLonP1) DJ-1 null MEFs growing under basal conditions and stained with Mitotracker (red), anti-LonP1 specific antibodies (green) and counterstained with DAPI (blue) for nuclei visualization. Scr DJ-1 null MEFs, panels C and E or sh mLonP1 DJ-1 null MEFs, panels D and F, were transiently transfected with hDJ-1 L10P, P158Δ and L166P (C and D) or hDJ-1 A107P, E163K and L172Q (E and F) and 48 hours after transfection were treated with 25µg/mL cycloheximide (CHX) for the times indicated. Panels show representative immunoblots with anti-DJ-1 polyclonal antibody of the indicated hDJ-1 missense mutants transfected MEFs. Anti-tubulin antibodies were used as total protein loading control. Quantifications are shown in the graphs below as mean ± s.e.m from three different experiments. Significant differences were found between scr and sh mLonP1 DJ-1 null MEFs transfected with P158Δ at the time points 2 hours (**p = 0.0001), 4 hours (**p = 0.003), 8 hours (**p = 0.0008) and 12 hours (**p = 0.009), between scr and sh mLonP1 DJ-1 null MEFs transfected with L166P at the time points 2 hours (**p = 0.0002), 4 hours (**p = 0.0001), 8 hours (**p = 0.0008) and 12 hours (**p = 0.002), between scr and sh mLonP1 DJ-1 null MEFs transfected with A107P at the time points 6 hours (**p = 0.0008), 12 hours (*p = 0.02) and 24 hours (**p = 8E-05), between scr and sh mLonP1 DJ-1 null MEFs transfected with E163K at the time points 12 hours (**p = 0.009) and 24 hours (*p = 0.01) and between scr and sh mLonP1 DJ-1 null MEFs transfected with L172Q at the time point 24 hours (**p = 0.001).

To ascertain no off-target effects of mLonP shRNA, human LonP1 (hLonP1) cDNA was used to rescue the phenotype of mLonP1 silenced cells respect to degradation of two of the unstable DJ-1 mutants.. As shown in Fig. 7, over expression of hLonP1 (Fig. 7A) completely rescue the inhibition of degradation of the unstable DJ-1 P158DELTA and L166P mutants produced by mLonP1 shRNAs. Taken together, those results clearly allowed the conclusion that LonP1 is the mitochondrial protease mainly responsible of the degradation of unstable DJ-1 mutants A107P, P158DELTA, E163K, L166P L172Q mutants. but not of the degradation of DJ-1 L10P mutant.

**Legend to Fig. 7.**
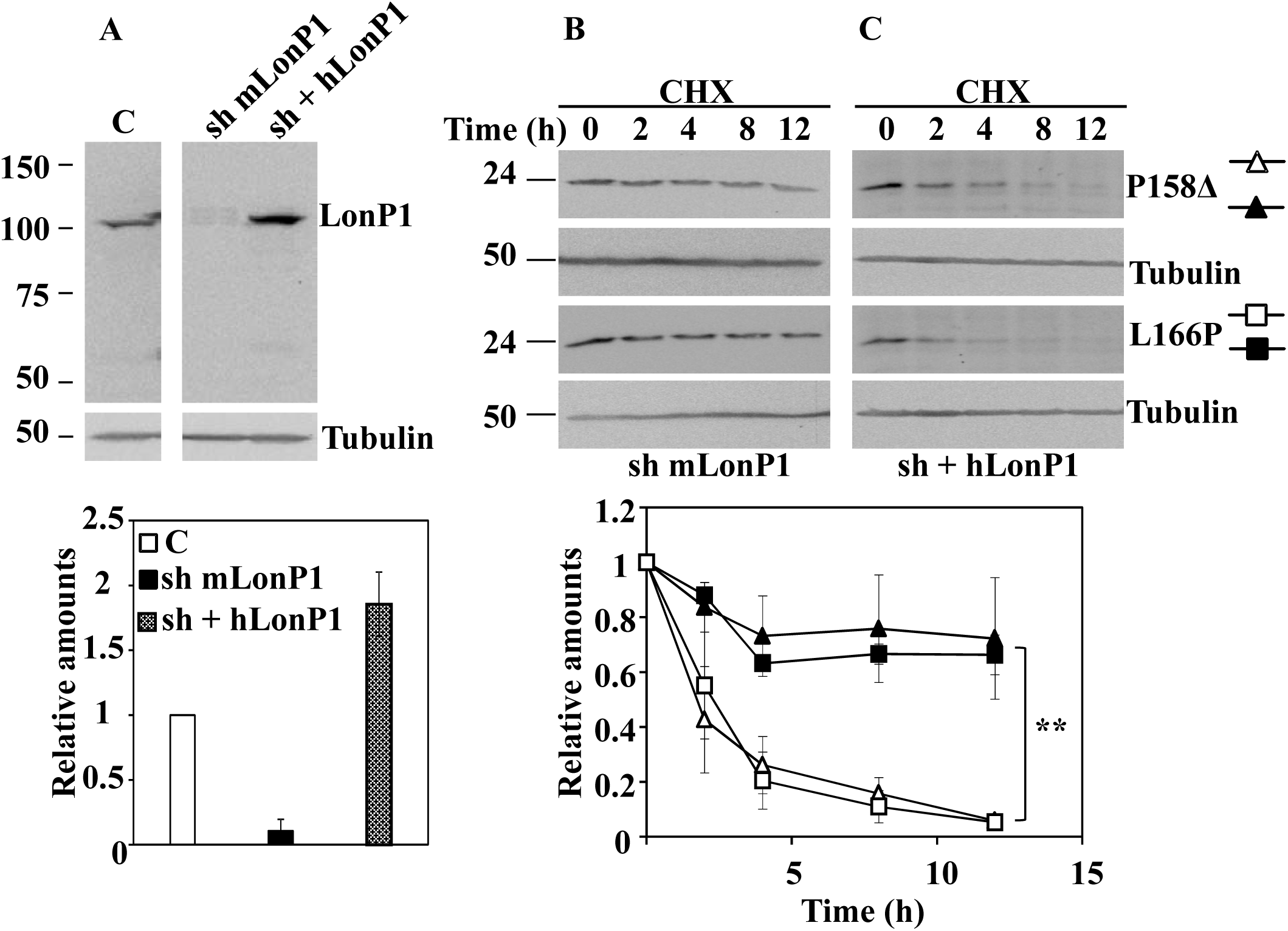
Rescue of the degradation of DJ-1 unstable mutants transfected in LonP1-silenced DJ-1 null MEFs by transfection of human LonP1. LonP1 shRNA-transduced DJ-1 null MEFs (sh mLonP1) were complemented with ectopically expressed human LonP1 (hLonP1). A, panel shows a representative immunoblot of LonP1 shRNA-mediated knockdown in DJ-1 null MEFs and the recovery of LonP1 protein levels after transfection with hLonP1. Immunoblot was developed with specific antibodies anti-LonP1. Quantification is shown in the right graph. LonP1-silenced DJ-1 null MEFs (sh mLonP1), panel B, and rescued LonP1-silenced DJ-1 null MEFs (sh + hLonP1), panel C, were transiently transfected with hDJ-1 P158Δ or L166P and 48 hours after transfection were treated with 25µg/mL cycloheximide (CHX) for the times indicated. Panels B and C show representative immunoblots developed with anti DJ-1 polyclonal antibody of the indicated constructs. Anti-tubulin antibodies were used as total protein loading control. Quantifications are shown in the graphs below as mean ± s.e.m from three different experiments. Significant differences were found between sh mLonP1 and sh + hLonP1 DJ-1 null MEFs transfected with P158Δ at the time points 2 hours (*p = 0.01), 4 hours (*p = 0.03), 8 hours (*p = 0.03) and 12 hours (*p = 0.04) and between sh mLonP1 and sh + hLonP1 DJ-1 null MEFs transfected with L166P at the time points 4 hours (*p = 0.01), 8 hours (**p = 0.001) and 12 hours (**p = 0.001).

## Discussion

DJ-1 WT is a stable protein when expressed by transfection in DJ-1 null MEFs, similar to what is found for endogenously expressed DJ-1 in other cells (Macedo et al., 2003) (Moore et al., 2003) (Schwanhausser et al., 2011) (Alvarez-Castelao et al., 2012a) (Song et al., 2016; Sanchez-Lanzas and Castano, 2018). The PD missense mutations A39S, E64D, A104T, D149A, K175E and the polymorphic missense variants R98Q and A171S are as stable as DJ-1WT, both in MEFs from DJ-1 null mice (Fig. 1) and in N2a cells, expressing the endogenous DJ-1 WT The DJ-1 point mutants L10P, A107P, P158DEL, E163K, L166P and L172Q were degraded with similar half-lives in MEFs from DJ-1 null mice and in N2a cells. The M26I DJ-1 mutant is slowly degraded in MEFs from null DJ-1,ice (see Fig. 1 and Supplementary Table 1), but degradation is not apparent in N2a cells after treatment of cells with CHX for 24h, as reported previously (Alvarez-Castelao et al., 2012a). These results indicated that the unstable (and untagged) DJ-1 mutants, qualitatively, are unstable even in the presence of the expression of endogenous DJ-1 WT. Several laboratories have reported (including ourselves)that the missense DJ-1 mutants are degraded by the ubiquitin-proteasome pathway: DJ-1 L166P (Bonifati et al., 2003) (Macedo et al., 2003) (Miller et al., 2003) (Moore et al., 2003) (Olzmann et al., 2004) (Takahashi-Niki et al., 2004) (Blackinton et al., 2005) (Gorner et al., 2007), (Alvarez-Castelao et al., 2012a), L10P and P158DEL (Ramsey and Giasson, 2010) (Rannikko et al., 2012) and L172Q (Taipa et al., 2016). Nevertheless, several groups reported that proteasome inhibitors are not able to completely prevent the degradation of the unstable mutants, L166P (Miller et al., 2003) (Olzmann et al., 2004) (Gorner et al., 2007) (Ramsey and Giasson, 2010), (Alvarez-Castelao et al., 2012a), L10P and P158DELTA (Ramsey and Giasson, 2010). Furthermore, as shown here, treatment with different inhibitors of the autophagic-lysosomal pathway (NH_4_Cl, leupeptin 3-methyl-adenine and E64) fail to prevent their degradation (Fig. 3). Similar results with those inhibitors have been reported previously for the degradation of DJ-1 L166P missense mutant (Miller et al., 2003) (Olzmann et al., 2004) (Blackinton et al., 2005).

In the search for alternative pathways of degradation, experiments of subcellular localization of DJ-1 by immunofluorescence and biochemical cell fractionation studies were performed (Figs. 4 & 5). Those data showed that the DJ-1 point mutants M26I, A107P, P158DEL, E163K, L166P and L172Q DJ-1, but not L10P, are significantly associated with mitochondria. Those results are in agreement with previously published work of the localization of M26I, P158DEL, E163K, L166P and L172Q (Bonifati et al., 2003) (Zhang et al., 2005) (Maita et al., 2013) (Bjorkblom et al., 2014a) (Kojima et al., 2016) (Taipa et al., 2016);, no data are available for A107P point mutant. The widely used experimental approach of using fusion constructs of the missense mutants with fluorescent proteins as reporters for both degradation and subcellular localization was discarded, as the DJ-1 L166P and M26I proteins fused to EGFP (Supplementary Fig, 3) did not meet the simple criteria of similar degradation rates and subcellular localization as the respective untagged DJ-1 missense mutants. Another clear demonstration of the caveat of using tagged version of proteins for studying DJ-1 point mutants. Similar results for the E63K mutant fused to GFP have been reported (Kojima et al., 2016).

To study the possible involvement of mitochondria in the degradation of missense DJ-1 mutants, we hypothesized that mitochondrial matrix LonP1 could be implicated. The experimental evidence presented (Fig. 6 and Supplementary Fig. 4) clearly indicated that mitochondrial LonP1 is implicated in the degradation of A107P, P158DEL, E163K, L166P and L172Q. We showed (Fig. 7) that the strong inhibition of their degradation by interrupting the expression of LonP1 can be rescued by overexpression of human LonP1, indicating the specificity of the observed inhibitory effects. In contrast, there is no effect on L10P degradation. the elucidation of the degradation pathway of this misesense mutant requires further investigation.

The localization of DJ-1 in the mitochondria and translocation to mitochondrial matrix have been studied (Maita et al., 2013) (Kojima et al., 2016). The L10P mutation located in strand 1 prevents homodimer formation while the mutant interacts with DJ-1 WT (Ramsey and Giasson, 2010) (Repici et al., 2013). The L10P mutation may hinder the mitochondrial localization, as that region of strand 1 of the 3D-protein structure (Honbou et al., 2003; Huai et al., 2003; Tao and Tong, 2003) has been implicated in mitochondrial localization. The DJ-1 E18A point mutant (a-helix 1) localizes to the mitochondria, but the substitution by alanine of the wild-type sequence at leucine (a.a. 7), valine (a.a. 8), isoleucine (a.a. 9) and leucine (a,a, 10, changed to Pro in the missense mutant) within strand 1 prevents mitochondrial localization and gives a nucleo-cytoplasmic subcellular distribution (Maita et al., 2013). The fact that the mutants C15DELTA and M26I (shown also here, Fig. 4) also localized to the mitochondria (Kojima et al., 2016), further supports the role of the aminoacid sequence of strand 1 and a-helix 1 (aminoacids 5 to 28) in the N-terminus as the sequence that either prevents the translocation of DJ-1 to mitochondria, or promotes its cytoplasmic retention. But those N-terminal structural elements are clearly insufficient to explain the location to mitochondria as DJ-1 mutants at the C-terminus that have an intact N-terminal sequence, also localize to the mitochondria. Clearly further experiments will allow to elucidate the structural requirements for the mitochondrial localization and matrix translocation of DJ-1 missense mutants.

Mutations in *LONP1* gene produce the CODAS syndrome with Cerebral, Ocular, Dental, Auricular and skeletal abnormalities (Dikoglu et al., 2015; Strauss et al., 2015), more recently a bi-allelic mutation (c.2282 C > T, (p.Pro761Leu) in *LONP1* results in neurodegeneration with deep hypotonia and muscle weakness, severe intellectual disability and progressive cerebellar atrophy (Nimmo et al., 2019). clearly showing a relationship of *LonP1* with central nervous system abnormalities and neurodegeneration. LonP1 is implicated in the degradation of matrix mitochondrial proteins like aconitase (Bota and Davies, 2002). 5-aminolevulinic acid synthase (Tian et al., 2011), TFAM (Lu et al., 2013), StAR (Granot et al., 2007), complex 1 of the OXPHOS (Pryde et al., 2016), SDH5 (Bezawork-Geleta et al., 2014) and SDHB (Maio et al., 2016) of complex II of OXPHOS and unfolded proteins. like the matrix located OTC Delta used to promote an unfolding protein response (mitUPR) in mitochondria (Jin and Youle, 2013; Bezawork-Geleta et al., 2015). LonP1 also participates in the default degradation in mitochondrial matrix of the PD linked protein kinase PINK1/PARK6. Under mitUPR conditions, PINK1 accumulates (increased accumulation was produced with silencing of LonP1) and recruits the PD-linked E3 ligase Parkin/PARK2 promoting mitochondrial clearance by mitoautophagy (Jin and Youle, 2013) (Greene et al., 2012). Other groups have observed the accumulation of PINK1 upon silencing LonP1 without the need of concomitant induction of mitUPR response (Thomas et al., 2014) (Zurita and Shoubridge, 2018). PINK1 is initially processed by the mitochondrial processing protease (MPP), other mitochondrial proteases like PARL, ClpPX and AFG3L2 also participate(Greene et al., 2012). The processed PINK1, in contrast to the above observations of degradation by LonP1, would be rapidly degraded by the ubiquitin-proteasome pathway (Lin and Kang, 2008; Narendra et al., 2010) after polyubiquitylation (Liu et al., 2017). In *Drosophila*, overexpression of DJ-1 rescues the altered phenotype caused by the loss of PINK1, but not of Parkin (Hao et al., 2010). Similarly, DJ-1 overexpression also rescues the increased sensitivity of *Substantia Nigra* to MPTP in PINK1 null mice (Haque et al., 2012). Furthermore, the mitochondrial fragmentation phenotype of DJ-1 null cells can be rescued by overexpression of PINK1 or PARKIN (Thomas et al., 2010) (Irrcher et al., 2010), Taken together all these results indicate that DJ-1 and PINK1 (PARKIN) probably acts in parallel pathways and LonP1 is involved in DJ-1 point mutants and PINK1 degradation, with variable degree of involvement of the ubiquitin-proteasome pathway.

In conclusion, the PD phenotype presented by patients harbouring homozygous mutations M26I, A107P, P158DEL, E163K, L166P and L172Q can be explained by a “loss of function”, similar to the effects produced by DJ-1 mutation that resulted in strong down-regulation or absence of DJ-1 mRNA (deletions, CNV and splicing mutations), because of its instability due to its mitochondrial targeting and degradation mainly by matrix mitochondrial LonP1 protease. A similar situation is the case of patients that are homozygous for DJ-1 L10P mutation, while its main pathway of degradation remains to be determined. In contrast, The other point mutants (A39S, E64D, A104T, D149A, K175E and A179T) and the rare polymorphisms (R98Q, A171S) could be that there are not truly pathogenic (likely sure for the polymorphic variants R98Q, A171S. Another possibility is that their half-life could still be lower than DJ-1 WT, but escape to our detection limits (24h), requiring the use of quantitative proteomic studies using SILAC pulse-chase experiments and MS for its determination (far away from the scope and budget of our work). By those methods the DJ-1 protein has a very long half-life (t1/2), from t1/2=187h in mouse NIH3T3 cells (Schwanhausser et al., 2011) to t1/2=199 to 299h as determine in vivo in mouse heart (Lau et al., 2016). Accordingly, the pathogenetic mechanism for A39S, E64D, R98Q, A104T, D149A, A171S, K175E and A179T remains to be determined.

## Materials and Methods

### Plasmid constructs

The vectors for expression of untagged human DJ-1 wild type (hDJ-1 WT) and missense mutants M26I, R98Q, A104T, D149A and L166P have been previously described (Alvarez-Castelao et al., 2012a). The mutant L10P was obtained by PCR (4 min, 97°C; 30 cycles of 45 s, 95°C; 1 min, 61°C and 1.5 min, 72 °C, and a final polymerization cycle of 15 min, 72°C) with the appropriate oligonucleotides, forward BamH1-hDJ-1 L10P: 5’ GAGCGGATCCATGGCTTCCAAGAGGGCTCTGGTCATCCCGGCTAAAGGAGCAG AGG 3’ and reverse XhoI-hDJ-1 L10P: 5’ GCGCCTCGAGCTAGTCTTTAAGAACAAGTGGAGCC 3’ and amplified DNA was subcloned into pCMV-Sport6 vector for expression in eukaryotes, as described (Alvarez-Castelao et al., 2012a).

The mutations A39S, E64D, A107P, P158Δ, E163K, A171S, K175E and A179T of hDJ-1 were introduced by PCR (initial 4 minutes, 97°C; 30 cycles of 45 seconds, 95°C; 1 minute, 61°C and 10 minutes, 72°C, and a final polymerization cycle of 15 minutes, 72°C) from hDJ-1 WT inserted in pCMV-Sport6 using the QuickChange™ Site-Directed Mutagenesis Kit (Statagene) with the following oligonucleotides: forward hDJ-1 A39S: 5’ CCGTTGCAGGCCTGTCTGGAAAAGACCCAGTAC 3’ and reverse hDJ-1 A39S 5’ GTACTGGGTCTTTTCCAGACAGGCCTGCAACGG 3’; forward hDJ-1 E64D: 5’ GCAAAAAAAGAGGGACCATTTGATGTGGTGGTTC 3’, and reverse hDJ-1 E64D: 5’ GAACCACCACATCAAATGGTCCCTCTTTTTTTGC 3’; forward hDJ-1 A107P: 5’ GATAGCCGCCATCTGTCCAGGTCCTACTGCTCTG 3’ and reverse hDJ-1 A107P: 5’ CAGAGCAGTAGGACCTGGACAGATGGCGGCTATC 3’; forward hDJ-1 P158Δ: 5’ CTGATTCTTACAAGCCGGGGTGGGACCAGCTTCGAG 3’ and reverse hDJ-1 P158Δ: 5’ CTCGAAGCTGGTCCCACCCCGGCTTGTAAGAATCAG 3’; forward hDJ-1 E163K: 5’ GGACCAGCTTTAAGTTTGCGCTTGC 3’ and reverse hDJ-1 E163K: 5’ GCAAGCGCAAACTTAAAGCTGGTCC 3’; forward hDJ-1 A171S 5’ GCAATTGTTGAATCCCTGAATGGCAAGGAGG 3’ and reverse hDJ-1 A171S: 5’ CCTCCTTGCCATTCAGGGATTCAACAATTGC 3’; forward hDJ-1 K175E: 5’ GCCCTGAATGGCGAGGAGGTGGCGGCTCAGG 3’ and reverse hDJ-1 K175E: 5’ CTTGAGCCGCCACCTCCTCGCCATTCAGGGC 3’; forward hDJ-1 A179T: 5’ GCAAGGAGGTGGCGACTCAAGTGAAGGC 3’ and reverse hDJ-1 A179T 5’ GCCTTCACTTGAGTCGCCACCTCCTTGC 3’. The hDJ-1 L172Q mutant was synthesized by GenScript.

The EGFP fusion protein constructs of hDJ-1 M26I and L166P were obtained by gene synthesis of hDJ-1 cDNA missense mutants including restriction sites 5’ for XhoI and 3’ for AgeI and subcloning into the XhoI-AgeI sites of the pEGFP-N1 mammalian expression vector (Genescript).

Mouse LonP1 shRNA (RMM4534-NM_028782) and the corresponding scramble control shRNA cloned in the lentiviral vector pLKO.1 puro (Quiros et al., 2014) were provided by Dr. Carlos López-Otín, Departamento de Bioquímica y Biología Molecular, Universidad de Oviedo, Spain. For rescue experiments in mouse LonP1 silenced cells, human LonP1 cDNA (clone MGC:1498 IMAGE:3350958) was obtained from the pOTB7 construct by digestion with EcoR1 and XhoI and subcloned into pCDNA 3.1 vector, previously digested with the same restriction enzymes. The DNA sequence of all constructs was verified by Sanger sequencing in an ABI Prism 3130XL.

### Cell culture, transfections and treatments

Mouse embryonic fibroblasts (MEFs) from null DJ-1 mice were provided by Dr. Jie Shen, Center for Neurologic Diseases, Brigham and Women’s Hospital and Harvard Medical School, Boston, USA (Goldberg et al., 2005) and grown in Dulbecco’s modified Eagle’s medium with low glucose (1 g/L) and supplemented with 15% fetal bovine serum and 50 µg/mL gentamicin. Null DJ-1 MEFs (3×10^6^) were transiently transfected with 6µg of the indicated DJ-1 constructs by nucleofection (Nucleofector II program T-016, Amaxa Nucleofector Technology) by using a in-house made transfection buffer (150mM KH_2_PO_4_, 24mM NaHCO_3_, 4mM D-glucose, 6mM ATP and 12mM MgCl_2_). Mouse neuroblastoma N2a cells were cultured in Dulbecco’s modified Eagle’s medium with 4.5 g/L glucose supplemented with 10% fetal bovine serum and 50 µg/mL gentamicin. N2a cells (3×10^5^) were transiently transfected using Lipofectamine 2000 reagent (Invitrogen) according to the manufacturer’s instructions.

Transfected cells were grown for 48h and then protein degradation was studied by treatment of cells with 25µg/mL cycloheximide (CHX) for the times indicated. At the dose of CHX indicated, cell viability (MEFs and N2a, trypan blue exclusion) was >95% after 24 hours of incubation. For some experiments, transiently transfected null DJ-1 MEFs were kept in complete medium or treated with 25µg/mL CHX in the absence or the presence of 10µM MG-132, 20mM NH_4_Cl, 20mM NH_4_Cl plus 50µM leupeptin (Leu) or 10 mM 3-methyl adenine (3MA) plus 5µM E64 for 12 or 24 hours, as indicated. Cells were then washed three times with cold phosphate-buffered saline (PBS), pelleted by centrifugation and used for the obtention of total cell extracts.

### Lentiviral production

For lentiviral production, 3.5×10^6^ HEK293T cells (cultured in Dulbecco’s modified Eagle’s medium with 4.5 g/L glucose supplemented with 10% fetal bovine serum and 50 µg/mL gentamicin) were seeded in 20mm x 100mm culture dishes and the next day transfected with 3.2µg of the packaging plasmid psPAX2, 1.6µg of the envelope plasmid pM2.G and 5µg of the corresponding lentiviral transfer plasmid, scramble control shRNA or mouse LonP1 shRNA in pLKO.1 puro plasmid (Quiros et al., 2014), by using Lipofectamine 2000 reagent according to the manufacturer’s protocol. The culture medium was removed 24 hours after transfection and complete fresh medium was added. Lentiviral particles were harvested 48 hours after transfection, again fresh culture medium was added and 72 hours after transfection a second harvest of lentiviral particles was collected. The collected culture media were centrifuged at 200 x g for 5 minutes, filtered-sterilized through 0.45µm filters (Millipore), aliquoted and stored frozen at -80°C until used for further experiments..

### Cell transduction for LonP1 silencing and rescue experiments

Null DJ-1 MEFs (1.3×10^5^ per well) or N2a cells (3×10^5^ per well) were seeded in 6-well culture plates and incubated with 700µL of filtered lentivirus-containing medium (scramble control shRNA or mouse LonP1 shRNA) in the presence of 8µg/mL polybrene to a final volume of 1.4mL per well. Culture media was replaced 24 hours after infection, 48 hours after infection transduced cells were selected by addition of 8µg/mL puromycin to the culture media for four days. Puromycin-resistant cells were transiently transfected with the indicated human DJ-1 constructs and treated with CHX for studying protein degradation, as described above.

For rescue experiments in LonP1 silenced MEFs, puromycin-resistant MEFs were transiently transfected with human LonP1, selected with culture medium containing 250µg/mL zeocin for four days, transiently transfected with the indicated human DJ-1 constructs and treated with CHX for studying protein degradation, as described above.

### Cell fractionation studies

For biochemical subcellular fractionation experiments, DJ-1 null MEFs were transfected with the indicated untagged human DJ-1 constructs and 48 hours after transfection cells were washed with cold PBS for three times, pelleted by centrifugation at 110 x g for 5 minutes at 4°C, suspended in lysis buffer (20mM HEPES pH 7.4, 250mM sucrose, 5mM MgCl_2_, 10mM KCl, 1mM EDTA, 1mM EGTA, 25mM NaF, 1mM orthovanadate, 10µM leupeptin, 1µg/mL pepstatin and 1mM PMSF) and incubated on ice for 10 minutes, with thoroughly pipetting up and down the suspension each 2 min. Next, the cell suspension was incubated for 5 minutes at room temperature with lysis buffer containing digitonin (to a final concentration of 50µg/mL), cell lysis was verified by Trypan blue staining. Cell suspensions were then transfered to ice and centrifuged at 1000 x g for 5 min at 4° C to remove nuclei, debris and non lysed cells. The supernatant was used as the total fraction (input) and was further centrifuged at 15000 x g for 15 minutes at 4°C. The supernatant was used as the cytosolic fraction whereas the pellet, containing mitochondria, was washed twice with 200µL of lysis buffer without digitonin and centrifuged at 15000 x g for 15 minute at 4°C. After the last wash, the pellet was resuspended in a final volume identical to the volume of the cytosolic fraction with a buffer containing 10mM HEPES pH 7.4, 10mM KCl, 1mM DTT, 0.6% NP-40, 1mM EDTA, 1mM EGTA, 25mM NaF, 1mM orthovanadate, 10µM leupeptin, 1µg/mL pepstatin and 1mM PMSF, vortexed for 10 seconds and centrifuged for 30 seconds at 15000 x g at 4°C. The supernatant of this centrifugation was used as the mitochondrial fraction. Samples from the total input, and mitochondrial and cytoplasmic fractions were loaded onto SDS-PAGE and analyzed by Western and immunoblot, as described below.

### Western immunoblotting

After the treatments, cells were directly lysed in SDS-buffer (62.5 mM Tris-HCl pH 6.8, 2% SDS, 20% glycerol, 10 µM leupeptin, 1 µg/mL pepstatin and 1 mM PMSF). Cell extracts were sonicated for 10 minutes on ice, centrifuged at 15000 x g for 10 minutes and supernatants used to measure total protein concentration with BCA protein assay kit (Thermo Scientific-Pierce, Waltham, MA, USA). Equal amounts of total protein were loaded onto 10% or 14% SDS-PAGE for Western transfer to PVDF membranes and immunoblotting.

Immunoblots were probed with mouse anti-DJ-1 monoclonal antibody (1:1,000, MBL. Clone 3E8); rabbit anti-DJ-1 polyclonal antibody (1:1,000, Abcam ab18257); rabbit anti-LonP1 polyclonal antibody (1:1,000, Abcam ab103809) and rabbit anti-Tim23 polyclonal antibody (1:1,000, Abcam ab230253). Mouse α-Tubulin (1:10,000, DM1A, Sigma) monoclonal antibody was used as loading control. The blots were developed with a peroxidase-labeled goat anti-mouse or anti-rabbit secondary antibody (1:5,000, Biorad) and chemiluminiscence detection (MF-ChemiBIS 3.2, DNR Bio-Imaging Systems). Blots were analyzed by quantitative densitometry using Totallab TL100 software (version 1.0, TotalLab Ltd., Newcastle upon Tyne, UK) and protein levels were normalized respect to tubulin.

### Immunofluorescence, confocal microscopy and image analysis

Cells grown on coverslips in 24-well plates were stained for 45 min with 250nM MitoTracker Red (Invitrogen). The coverslips were then washed with cold PBS three times, fixed with 4% paraformaldehyde in PBS for 20 min at room temperature, permeabilized in PBS with 0.1% Triton X-100 and blocked with PBS and 3% BSA for 1h at room temperature. The coverslips were processed for indirect immunofluorescence by incubation with rabbit anti-DJ-1 polyclonal antibody (1:200, Abcam) or rabbit anti-LonP1 polyclonal antibody (1:100, Abcam) for 3 hours at room temperature, washed with PBS three times for 10 min followed by incubation with Alexa-488 or Alexa-647 fluorescent-labelled secondary antibodies (1:1,000) in blocking buffer for 1 hour. Next, coverslips were washed with PBS three times for 10 minutes and 1 μg/mL DAPI (4’,6-diamidino-2-phenylindole) was included for nuclear counterstaining in the second of the washing steps. Coverslips with transfected cells with EGFP fusion protein constructs of DJ-1 were processed for direct fluorescence visualization by incubation with 1 μg/mL DAPI for 5 minutes. Coverslips were mounted with Prolong Gold antifade reagent (Invitrogen) for confocal microscopy observation in a laser scanning microscope (Leica TCS SP5).

To quantify the co-localization between MitoTracker fluorescence (red channel) and the immunofluorescence of transfected human DJ-1 constructs (green channel), the confocal fluorescence images obtained were analysed with the Image Correlation Analysis (ICA) plugin of ImageJ software (Schneider et al., 2012). After background subtraction, a single cell was defined as a region of interest (ROI) and quantitative co-localization analysis of the pixels for both channels (red and green) was measured using Pearson’s correlation coefficient the hDJ-1 / MitoTracker. Data are presented as mean ± s.e.m from at least 20 individual cells from two different experiments.

### Statistical Analysis

Quantitative data are reported as means ± s.e.m from three different experiments. When indicated, values are expressed as mean ± upper and lower values from two different experiments. Statistical significance between groups was calculated using a two-tailored Student’s t-test.

## Acknowledgments

This work was supported by grants from MICIU SAF2017-85199, and by grant S2017/BMD-3700 (NEUROMETAB-CM) from Comunidad de Madrid co-financed with the Structural Funds of the European Union.to JGC. We thanks Dr. Carlos López-Otín, Departamento de Bioquímica y Biología Molecular, Universidad de Oviedo, Spain for providing us the shRNA constructs for LonP1 used in this work.

---

**Supplementary Table 1.**
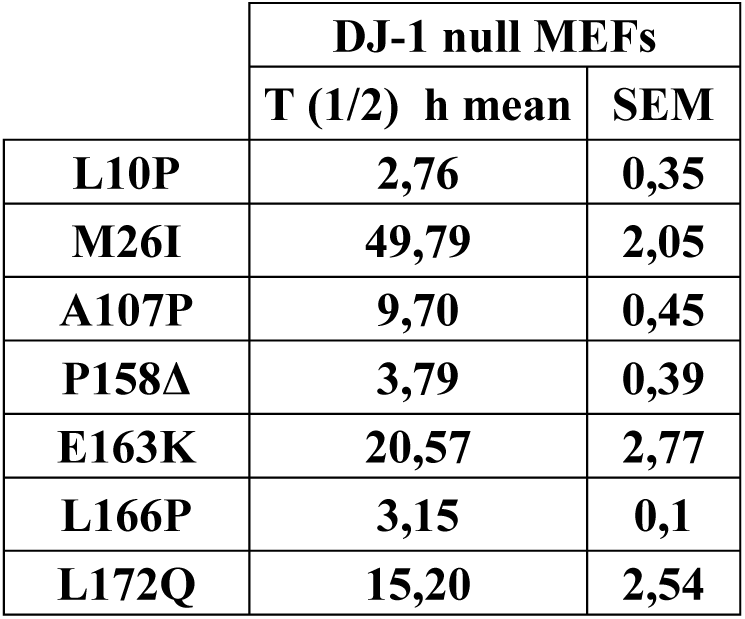
Half-life of unstable human DJ-1 missense mutants transfected in DJ-1 null MEFs. Reported values of the half-life (T ½) of these mutants in DJ-1 null MEFs are from the data of the experiments presented in Fig. 1. Values are expressed in hours (h) as mean ± s.e.m. from three different experiments.

## Legends to Supplementary Figures

**Legend to Supplementary Fig. 1.**
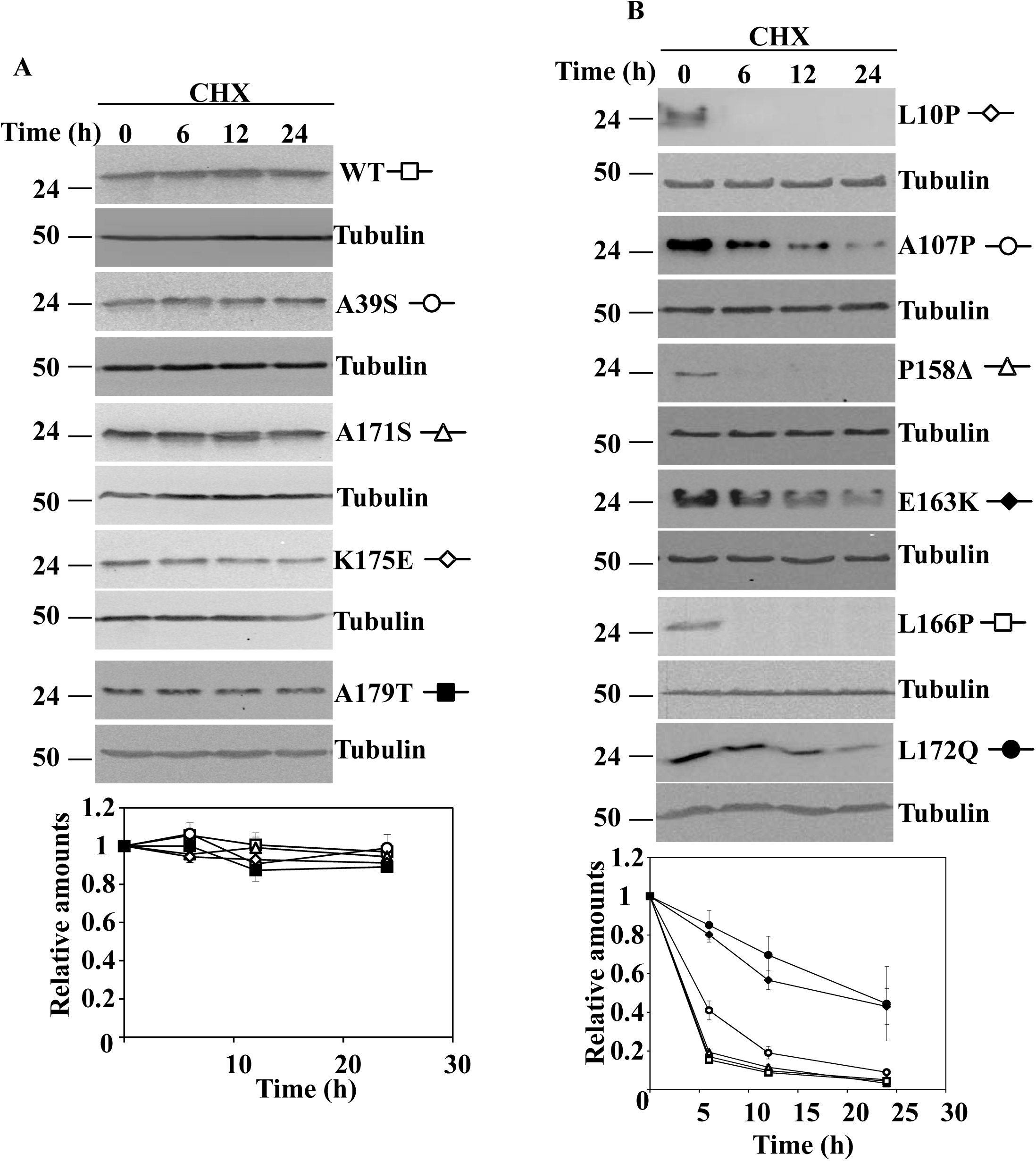
Degradation of human DJ-1 wild type and missense mutants in N2a transfected cells. N2a cells were transiently transfected with the indicated untagged human DJ-1 (hDJ-1) constructs and 48h after transfection were treated with 25µg/mL cycloheximide (CHX) for the times indicated. A, panels show representative immunoblots with anti-DJ-1 monoclonal antibody of hDJ-1 wild type (WT), A39S, A171S, K175E and A179T transfected cells. B, panels show representative immunoblots with anti-DJ-1 monoclonal antibody of hDJ-1 L10P, A107P, P158Δ, E163K, L166P and L172Q transfected N2a cells. Anti-tubulin antibodies were used as total protein loading control. Below are shown the graphs of quantification of the corresponding immunoblots. Data are mean ± s.e.m from three different experiments.

**Legend to Supplementary Fig. 2.**
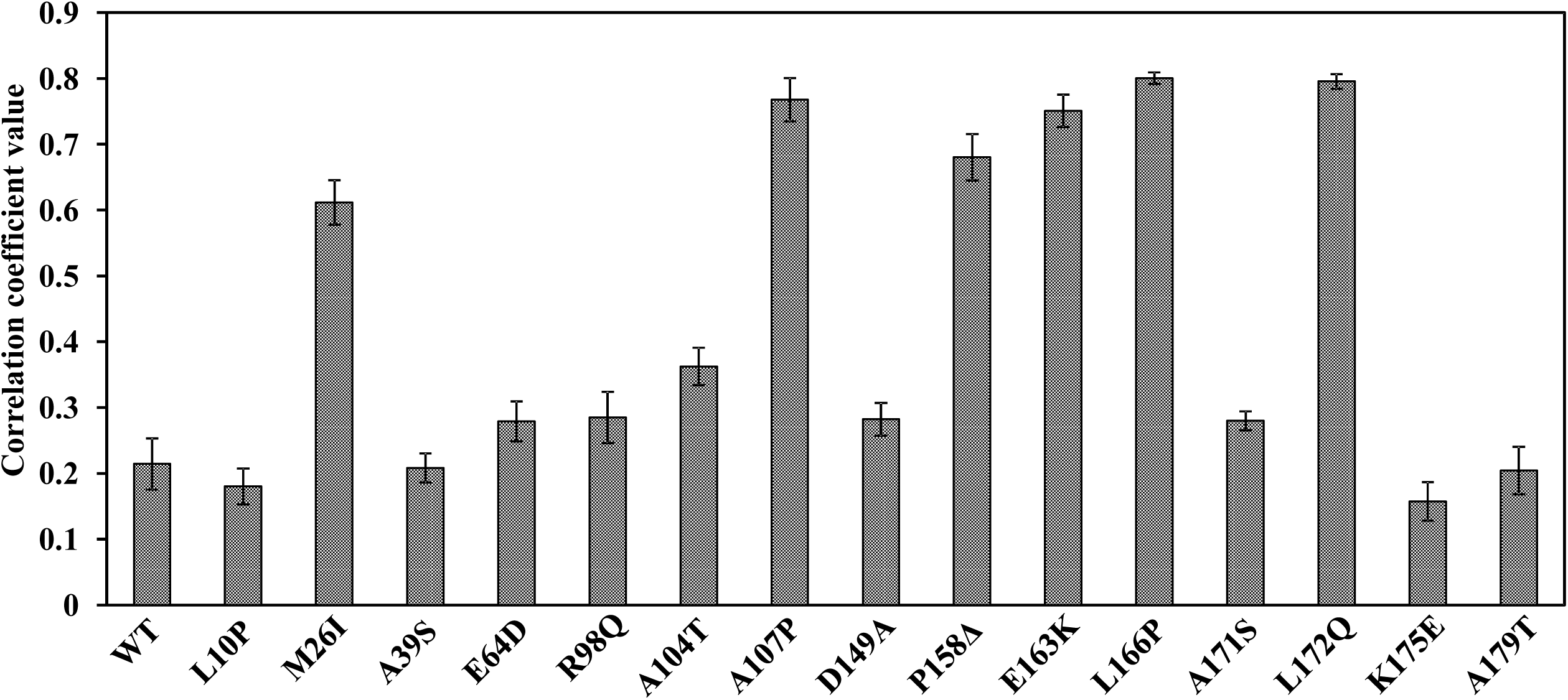
Analysis of the co-localization of Mitotracker fluorescence and immunofluorescence of DJ-1 wild type and missense mutants in transfected DJ-1 null MEFs. DJ-1 null MEFs were transfected with the indicated untagged human DJ-1 constructs, stained with Mitotracker (red channel) and processed for immunofluorescence with anti-DJ-1 (green channel) polyclonal specific antibodies. Graph shows the quantitative co-localization analysis between DJ-1 and Mitotracker performed in the confocal fluorescence images obtained as measured by the mean of the DJ-1 / Mitotracker Pearson’s correlation coefficient. Data are presented as mean ± s.e.m from at least 20 individual cells.

**Legend to Supplementary Fig. 3.**
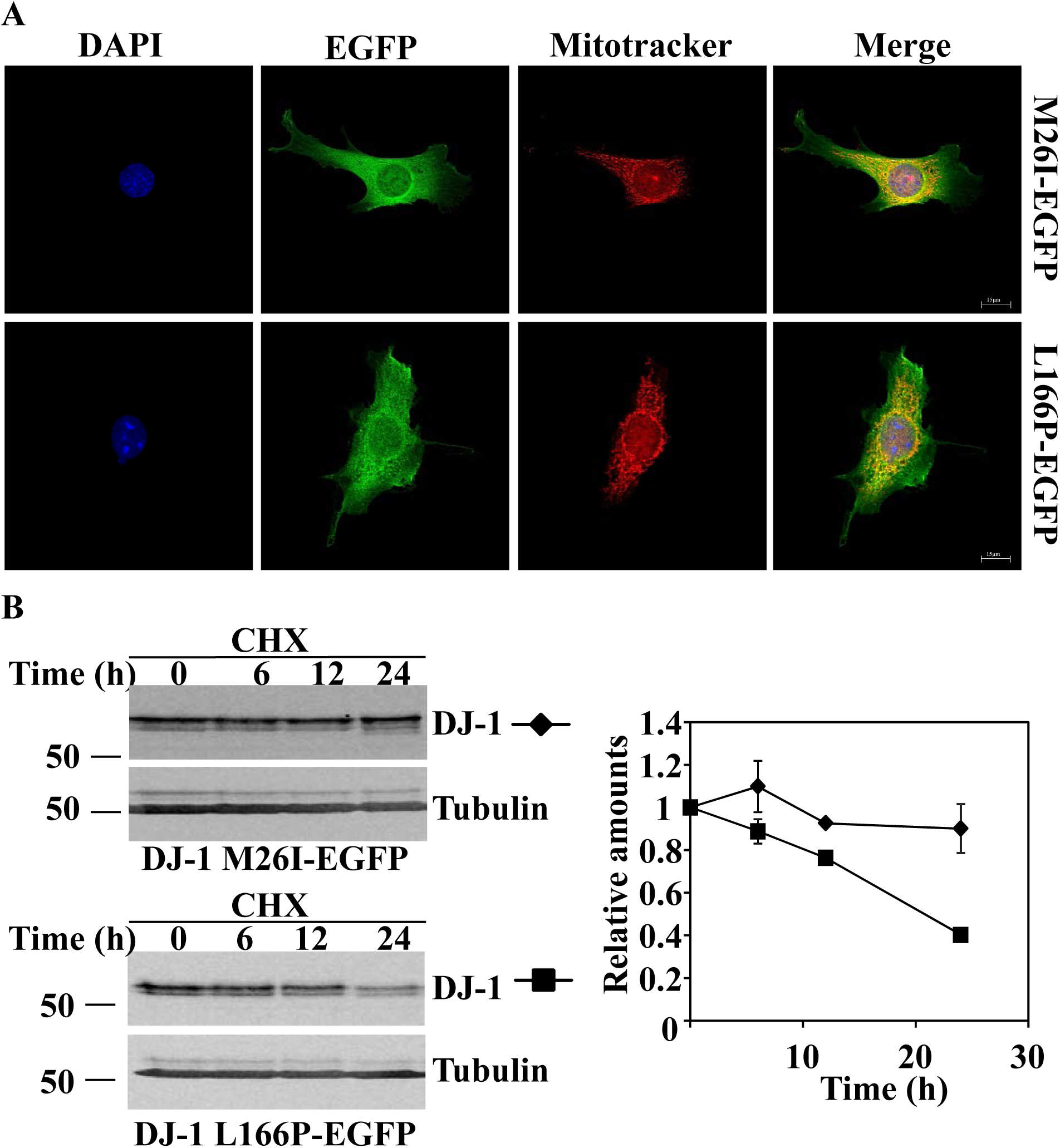
Subcellular localization by direct fluorescence and degradation of M26I and L166P DJ-1 - EGFP fusion constructs transfected in DJ-1 null MEFs. A, confocal fluorescence localization of M26I-EGFP or L166P-EGFP in transfected DJ-1 null MEFs growing under basal conditions, stained with Mitotracker (red), analysed by direct fluorescence of EGFP fusion protein (green) and counterstained for nuclei visualization with DAPI (blue). M26I-EGFP or L166P-EGFP transfected DJ-1 null MEFs were treated with 25µg/mL cycloheximide (CHX) for the times indicated and total cell lysates were analysed by Western and immunoblot. B, panels show representative immunoblots with anti-DJ-1 polyclonal antibody of M26I-EGFP and L166P-EGFP transfected cells. Anti-tubulin antibodies were used as total protein loading controls. Quantification of the corresponding immunoblots is shown in the graphs of the right. Data are expressed as mean ± upper and lower limit from two different experiments.

**Legend to Supplementary Fig. 4.**
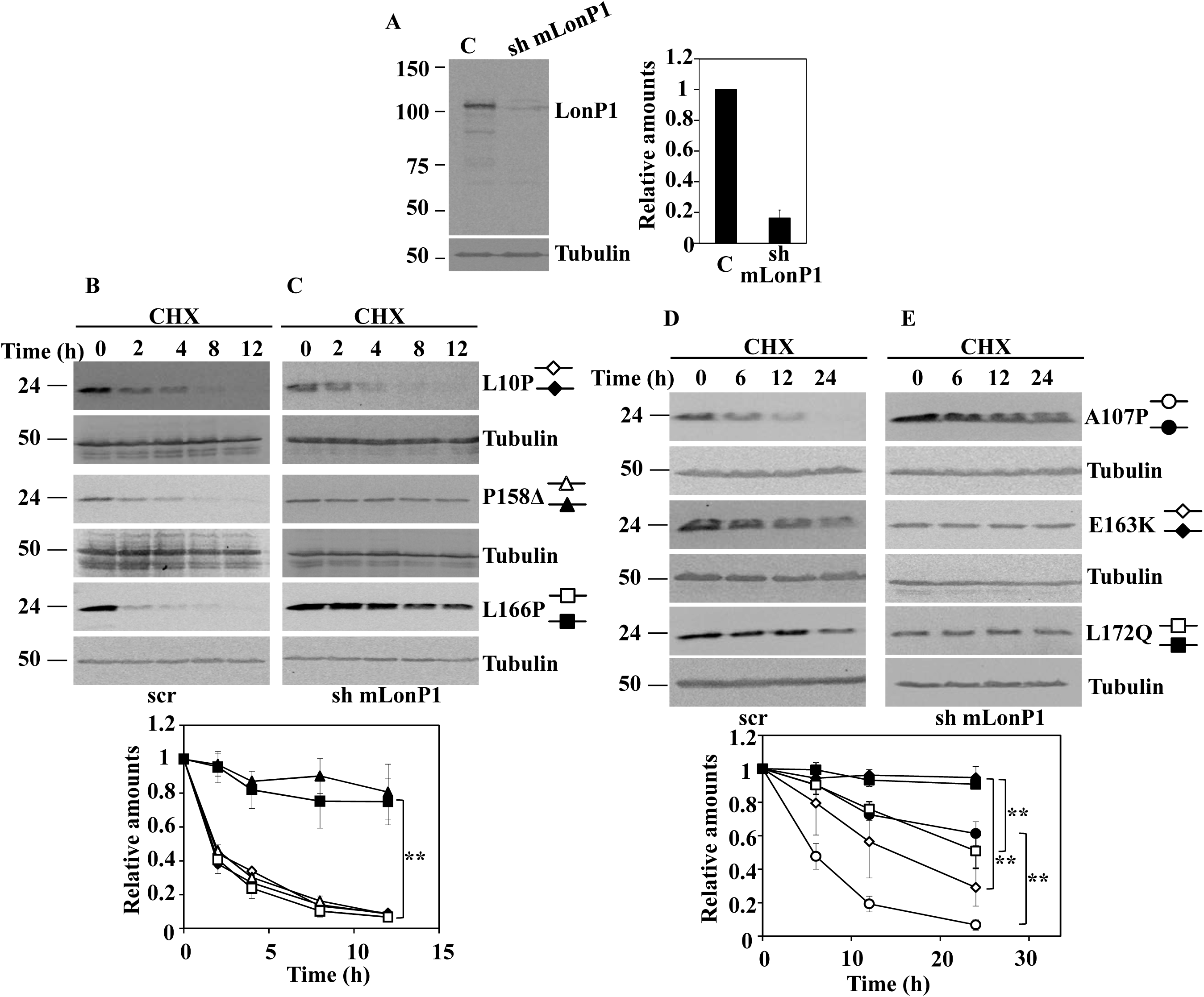
Effect of LonP silencing on the degradation of unstable DJ-1 mutants transfected in N2a cells. N2a cells were transduced with either non-target shRNA (scr) or LonP1 shRNA (sh mLonP1) as described under the material and method section. A, panel shows a representative immunoblot of LonP1 shRNA-mediated knockdown in N2a cells developed with specific antibodies anti-LonP1. Quantification is shown in the right graph. B, Non-target shRNA lentiviral transduced (scr) or LonP1 shRNA lentiviral transduced (sh mLonP1) N2a cells were transiently transfected with the indicated untagged hDJ-1 constructs and 48 hours after transfection were treated with 25µg/mL cycloheximide (CHX) for the times indicated. Panels B (scr) and C (sh mLonP1) show representative immunoblots with anti-DJ-1 monoclonal antibody of hDJ-1 L10P, P158Δ and L166P transfected N2a cells. Panels D (scr) and E (sh mLonP1) show representative immunoblots with anti-DJ-1 monoclonal antibody of hDJ-1 A107P and E163K transfected cells. Anti-tubulin antibodies were used as total protein loading control. Quantifications are shown in the graphs below as mean ± s.e.m from three different experiments. Significant differences were found between scr and sh mLonP1 N2a cells transfected with P158Δ at the time points 2 hours (**p = 0.002), 4 hours (**p = 3.4E-07), 8 hours (**p = 0.002) and 12 hours (*p = 0.01), between scr and sh mLonP1 N2a cells transfected with L166P at the time points 2 hours (**p = 0.006), 4 hours (**p = 0.009), 8 hours (*p = 0.01) and 12 hours (**p = 0.008), between scr and sh mLonP1 N2a cells transfected with A107P at the time points 6 hours (p = 0.03), 12 hours (**p = 0.0008) and 24 hours (**p = 0.002), between scr and sh mLonP1 N2a cells transfected with E163K at the time points 12 hours (*p = 0.04) and 24 hours (**p = 0.005) and between scr and sh mLonP1 N2a cells transfected with L172Q at the time points 12 hours (*p = 0.04) and 24 hours (*p = 0.01).

